# Vinculin mediated axon growth requires interaction with actin but not talin

**DOI:** 10.1101/2020.06.29.177758

**Authors:** Pranay Mandal, Vivek Belapurkar, Deepak Nair, Narendrakumar Ramanan

## Abstract

Axon growth requires coordination of the actin cytoskeleton by actin-binding proteins in the extending neurites. Vinculin is a major constituent of focal adhesion but its role in neuronal migration and axon growth is poorly understood. We found that vinculin deletion in mouse neocortical neurons attenuated axon growth both *in vitro* and *in vivo*. Using different functional mutants of vinculin, we found that expression of a constitutively active vinculin significantly enhanced axon growth while the head-neck domain had a moderate inhibitory effect. Interesting, we found that vinculin-talin interaction was dispensable for axon growth and neuronal migration. Strikingly, expression of the tail domain delayed migration, increased branching and stunted axon. Inhibition of the Arp2/3 complex or abolishing the tail domain interaction with actin completely reversed the branching phenotype caused by tail domain expression without affecting axon length. Super-resolution microscopy showed increased mobile fraction of actin in tail domain expressing neurons. Our results provide novel insights into the role of vinculin and its functional domains in regulating neuronal migration and axon growth.

## Introduction

Neuronal migration, axon growth and extension are intricately regulated processes that must be strictly coordinated for normal structural and functional development of the mammalian brain. These require coordination of the cellular cytoskeleton at the leading processes during neuronal migration and at the growth cone during axon growth (Dent *et al*, 2011; Lian & Sheen, 2015). The growth cone at the axon tip is a highly dynamic structure that is in constant interaction with the environment. The growth cone movements are aided by the formation of actin-rich membranous structures called lamellipodia and filopodia (Luo, 2002). These structures are driven by constant polymerization and depolymerization of the actin cytoskeleton. The actin cytoskeletal dynamics are regulated by several actin-binding proteins (Ishikawa & Kohama, 2007), which mediate polymerization, depolymerization and branching of actin filaments.

A critical first step in the growth and extension of neurites is filopodia formation. Filopodia are F-actin rich structures that are essential for neurite formation and growth (Dent *et al*, 2007). Studies have shown that neurons that lack critical actin-binding proteins showed attenuated neuritogenesis and axon growth (Kwiatkowski *et al*, 2007). Extension of the cell membrane and growth of axon requires constant engagement of the actin cytoskeleton with the extracellular matrix (ECM) through integrin receptors on the cell surface (Letourneau *et al*, 1988). This engagement and the force required to push the membrane forward is generated by vinculin and talin (Mierke, 2009; Romero *et al*, 2020). Talin binds the integrin receptors and activates them and promotes their interaction with ECM (Calderwood *et al*, 2002; Garcia-Alvarez *et al*, 2003). Vinculin is then recruited to focal adhesion sites and serves as a link between Talin/integrin receptors and the actin cytoskeleton (Atherton *et al*, 2015; Romero *et al*., 2020).

Vinculin is a 117 kDa protein made up of three major domains – an N-terminal head (Vh), polyproline linker and a C-terminal tail domain (Vt) (Ziegler *et al*, 2006). Vinculin exists in an auto-inhibited state in which all the ligand-binding sites are masked by intramolecular interactions between the head and tail domains (Bakolitsa *et al*, 2004; Borgon *et al*, 2004; Johnson & Craig, 1994, 1995). Vinculin activation results in conformation changes that disassociates head-tail interaction exposing the binding sites (Bakolitsa *et al*., 2004; Borgon *et al*., 2004). Vinculin is found in neuronal growth cones and binds to talin and F-actin through its head and tail domains respectively and to several other actin-binding proteins (Burridge & Mangeat, 1984; Letourneau & Shattuck, 1989; Menkel *et al*, 1994; Ziegler *et al*., 2006). Vinculin has been shown to play a role in cell adhesion and motility in fibroblasts and PC12 cells. Vinculin null mice exhibit embryonic lethality at E10 with severe defects in heart and neural tube development (Xu *et al*, 1998a). Vinculin-deficient fibroblasts showed poor adhesion to substrate, less spreading, enhanced cell migration and cytoskeletal dynamics (Coll *et al*, 1995; Demali, 2004; Thievessen *et al*, 2013; Xu *et al*, 1998b). Vinculin has been shown to be important for filopodia and lamellipodia stability, neurite outgrowth and axon growth *in vitro* (Li *et al*, 2014; Sydor *et al*, 1996; Varnum-Finney & Reichardt, 1994). Despite these studies, the requirement for vinculin and the role of different functional domains of vinculin for neuronal migration and axon growth *in vivo* is poorly understood.

In this study, we show that vinculin deletion in cortical neurons affects neuronal migration and attenuates axon growth both *in vitro* and *in vivo*. Furthermore, we found that different domains of vinculin affect axon growth to different extents. Expression of a constitutively active vinculin significantly enhanced axon growth while the tail domain alone attenuated axon growth. Vinculin tail domain expression caused an abnormal and enlarged cell soma, and highly branched neurites and stunted axon. Interestingly, abolishing vinculin-talin interaction did not affect neuronal migration and axon growth. The vinculin tail-induced branching phenotype was dependent on F-actin and Arp2/3 complex interactions. Thus, our findings provide novel insights into the role of vinculin in neuronal migration and axon growth.

## Results and Discussion

### Vinculin is necessary for axon growth

To understand the requirement of vinculin for axon growth, we first ablated vinculin in neocortical neurons using CRISPR/Cas9-mediated gene deletion (Suppl. Fig. 1A, B). Neocortical neurons from postnatal day 1 (P1) mice were transfected either with empty Cas9n-T2A-GFP (Cas9-nickase) expressing plasmid or Cas9n-T2A-GFP plasmid along with a pair of guide RNAs (gRNAs) targeting exon 1 of *Vcl* gene. Neurons were fixed at 4 days post electroporation and plating and immunostained for β-III tubulin to label all neurites and GFP to identify transfected neurons. The control plasmid expressing neurons exhibited normal axon growth. In contrast, neurons expressing gRNAs targeting *Vcl* showed highly attenuated axon growth (Suppl. Fig. 1C, D). This was consistent with an earlier finding in which shRNA-mediated knockdown of vinculin was shown to attenuate axon growth in cultured hippocampal neurons (Li *et al*., 2014). Next, we asked whether vinculin deletion also affected axon growth *in vivo*. For this, we introduced vinculin gRNAs into neural progenitor cells by *in utero* electroporation in embryonic day 14.5 (E14.5) embryos and analysed axon growth at E17.5. In the empty Cas9n-GFP plasmid (control) transfected embryos, the GFP^+^ cells in the neocortex showed longer axons. However, the vinculin gRNA expressing neurons had much shorter axons (Suppl. Fig. 1E, F) suggesting that vinculin is necessary for axon growth. Vinculin null fibroblasts have been shown to exhibit increased cell mobility in cultures (Coll *et al*., 1995; Xu *et al*., 1998b). We therefore analysed neuronal migration at E17.5 following vinculin deletion in E14.5 embryos. In control plasmid electroporated embryos, we found GFP^+^ cells present in all the layers of the neocortex (Suppl. Fig. 1G, H). In contrast, most of vinculin gRNA expressing neurons had migrated to the superficial layers indicating faster migration rate (Suppl. Fig 1G, H). Furthermore, we found that the vinculin gRNA expressing neurons exhibited shorter leading process compared to the control GFP^+^ neurons (Suppl. Fig 1I, J). Together, these findings suggest that vinculin deletion attenuates neurite growth and enhances cell migration *in vivo*.

### Vinculin functional domains affect axon growth and cell migration differently

Vinculin has three major domains – head, neck and tail domains, through which it interacts with other actin-binding proteins, kinases and F-actin. But the requirements for these domains for axon growth and neuronal migration is poorly understood. To study this, we expressed several N-terminal GFP-tagged vinculin mutants in cultured neocortical neurons and assessed for axon growth at 4 days *in vitro* (DIV) (Cohen *et al*, 2005; Cohen *et al*, 2006; Humphries *et al*, 2007) (Suppl. Fig. 2). These mutants included vinculin full length (Vcl-FL), vinculin N-terminal domain lacking the tail domain (Vcl-258, amino acids 1-258; Vcl-851, amino acids 1-851), vinculin tail domain (amino acids 881-1066) that binds actin (Vcl-T), a constitutively active form of vinculin (Vcl-T12) that has point mutations abolishing the head-tail interactions and a vinculin mutant that cannot bind PIP2 (Vcl-LD) (Chandrasekar *et al*, 2005; Cohen *et al*., 2005). Expression of each of these mutants affected axon growth to varying extents (Fig. 1A, B). Vcl-FL expression did not increase axon growth any further compared to control plasmid transfected cells. However, Vcl-T12 expression significantly increased axon growth while the other mutants attenuated axon growth (Fig. 1B). We next asked whether Vcl-T12 also increased axon growth *in vivo*. For this, we electroporated embryos at E14.5 and analysed axon growth at P5. For comparison, we chose Vcl-T, which caused a severe attenuation of axon growth in cultured neurons. Similar to that seen in cultured cells, Vcl-T12 was able to increase axon growth *in vivo* compared to control GFP^+^ neurons. In contrast, Vcl-T neurons exhibited highly stunted axon growth (Fig. 1C, D). Together these findings suggest that the full-length vinculin is required for normal axon growth while a constitutively active vinculin, that exists in an open conformation, can augment axon growth.

**Figure 1.**
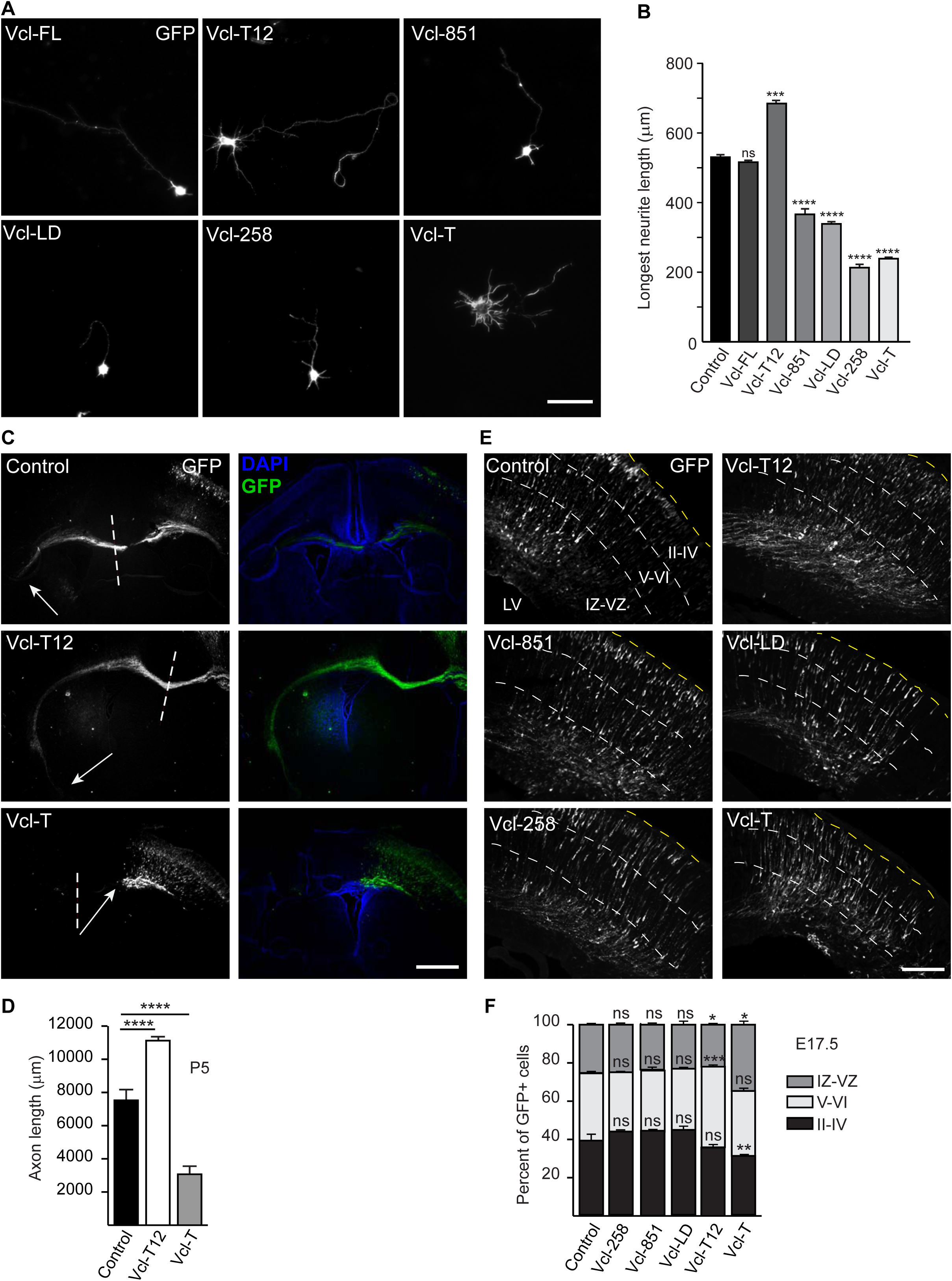
Different requirement of functional domains of vinculin for axon growth and cell migration *in vivo* and *in vitro*. **(A)** Primary neocortical neurons from P0.5 mouse pups were electroporated with different N-terminal GFP-tagged domain constructs of *Vinculin* and were grown for 4 DIV and immunostained for GFP. Absence of either head and neck (Vcl-T), neck and tail (Vcl-258) or tail alone (Vcl-851) and PIP2-binding defective mutant (Vcl-LD) attenuated axonal growth, while the constitutively active vinculin (Vcl-T12) enhanced axon growth while the full-length (Vcl-FL) did not have any effect. **(B)** Quantification of the length of longest neurite of neurons expressing different mutants of vinculin shown in (A) (n=250-300 cells). **(C)** Neocortical progenitors in E14.5 embryos were transfected with the domain deleted constructs of vinculin by *in utero* electroporation (IUE). The pups were sacrificed at P5 and axons were visualized using anti-GFP immunostaining (green). The Vcl-T transfected neurons exhibited highly stunted axons compared to the control GFP neurons. In contrast, Vcl-T12 expression increased axon growth significantly. Arrows indicate the axon terminal along the corpus callosum. Dashed lines represent the longitudinal fissure separating the two cerebral hemispheres. **(D)** Quantification of axon length from (C) (n=3 animals). **(E)** IUE was performed at E14.5 and the embryos were harvested at E17.5. 20µm brain sections were prepared and immunostained for GFP to study cell migration. The number of GFP^+^ neurons in different cortical layers were counted. LV, lateral ventricle; IZ/VZ, intermediate zone/ventricular zone; V-VI, layers 5/6; II-IV, layers 2-4. **(F)** Quantification of the percentage of GFP^+^ neurons in different layers of cortex from (E). More Vcl-T expressing neurons were seen in the lower layers compared to control indicating delayed migration whereas Vcl-T12^+^ neurons showed faster migration compared to control. (\**p*=0.0254, Ctrl-Vcl-T12, \**p*=0.0258, Ctrl-Vcl-T; n=3 animals). Scale, 100µm (A), 2mm (C), 400µm (E). \*\**p*<0.01, \*\*\**p*<0.001, \*\*\*\**p*<0.0001. ns, non-significant. One-way ANOVA and Tukey’s *post-hoc* test in (B) and (F). Two-tailed student’s *t* test in (D).

We next asked about the requirement for these domains for neuronal migration. During corticogenesis, neurons generated in the ventricular/subventricular zone migrate radially towards the pial surface and form distinct layers in an inside-out manner termed neocortical lamination (Nadarajah & Parnavelas, 2002). We transfected the above vinculin mutants in E14.5 embryos and assessed cell migration three days later at E17.5. We did not see any defects in cell migration when Vcl-258 and Vcl-851 were expressed as compared to control GFP transfection (Fig. 1E, F). However, when the constitutively active Vcl-T12 mutant was expressed, more GFP^+^ cells were seen in layer 5/6 and less number in the ventricular zone/intermediate zone (VZ/IZ). But there was no change in the number of cells in the upper layers 2/3. In contrast, Vcl-T expressing cells showed delayed migration to the upper layers and an increase in the number of cells in the ventricular zone (Fig. 1E). Also, there were fewer cells in layers 2/3. Together these findings indicated that lack of the vinculin tail domain (Vcl-258 and Vcl-851) does not affect neuronal migration while vinculin tail domain expression severely delayed cell migration from the ventricular zone/intermediate zone. Vcl-T12 appears to enhance migration from the intermediate zone compared to Vcl-T mutant, but not to the extent seen with vinculin deletion (Fig 1E, F). Given this interesting phenotype observed with Vcl-T12 and Vcl-T mutants, we next asked how these mutants affected migration at a later time point. For this, we electroporated embryos at E14.5 and analysed cell migration at P5. The Vcl-T12 expressing neurons that were seen enriched in layers 5/6 have reached their upper layer targets and there was no significant difference compared to control GFP expressing cells. In striking contrast, a greater number of Vcl-T expressing neurons were still seen in the lower layers (VZ/IZ and layers 5/6) (Fig. 2A, B). We further confirmed the defect in neuronal migration using immunostaining for Cux1, which is specifically expressed in layers 2-4 in the neocortex (Ferrere *et al*, 2006). We observed more Cux1^+^ cells in the lower layers in Vcl-T transfected embryos than those expressing control GFP (Suppl. Fig. 3). Interestingly, while analysing cell migration, we noticed that Vcl-T expressing neurons exhibited abnormal cell soma morphology. We analysed the area of neuronal soma and found that Vcl-T expressing cells had a much larger cell soma compared to control neurons (Fig. 2C, D).

**Figure 2.**
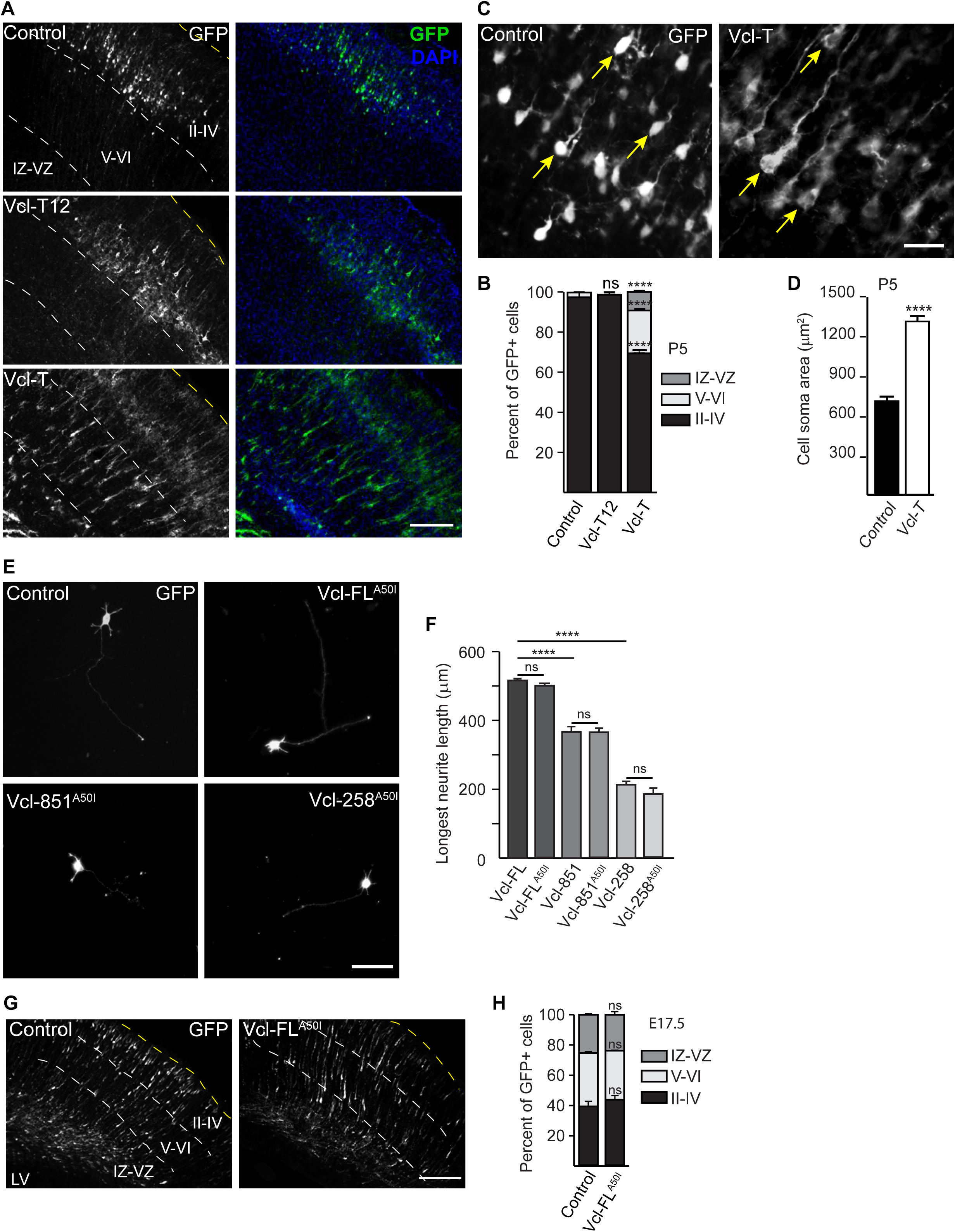
Interaction of talin with vinculin is dispensable for axonal growth and neuronal migration. **(A)** Neocortical progenitors in E14.5 embryos were electroporated by IUE with control GFP, GFP-Vcl-T12 or GFP-Vcl-T and pups were analysed at P5 for neuronal migration. Vcl-T expressing neurons were still in migratory phase even at P5 while Vcl-T12 expressing cells had migrated to upper layers of cortex. **(B)** Quantification of the percentage of GFP^+^ neurons in different layers of cortex from (A) (n=3 animals). **(C)** Representative images of neurons transfected with either empty GFP (control) or Vcl-T *in vivo* in E14.5 embryos. The Vcl-T expressing neurons exhibited larger and abnormal cell soma compared to control neurons. **(D)** Quantification of cell soma area from (C) (n=100 cells, n=3 animals). **(E)** Primary mouse neocortical neurons were electroporated with GFP-tagged vinculin constructs that contained the head domain but were defective in talin binding (Vcl-FL^A50I^, Vcl-258^A50I^ and Vcl-851^A50I^). Length of the longest neurite was compared with the corresponding mutant having intact talin binding site (Vcl-FL, Vcl-258 and Vcl-851). **(F)** Quantification of longest neurite length from (E). Abolishing talin-vinculin interaction does not affect axonal growth *in vitro*. (n=3 animals). **(G)** IUE was performed using control GFP and Vcl-FL^A50I^ plasmids and the embryos were harvested at E17.5 to study migration. Inhibition of talin-vinculin interaction did not affect neuronal migration. **(H)** Quantification of the percentage of GFP-positive neurons in different layers of cortex (G). (n=3 animals) (LV, lateral ventricle; IZ/VZ, intermediate zone/ventricular zone; V-VI, layers 5/6; II-IV, layers 2-4). Scale, 400µm (A, G), 150µm (C), 100µm (E). \*\*\*\**p*<0.0001; ns, non-significant. One-way ANOVA and Tukey’s *post-hoc* test (B, F and H). Two-tailed student’s *t* test (D).

### Talin binding is dispensable for vinculin-dependent axon growth

Previous studies in fibroblasts have shown the interaction of vinculin with talin, a focal adhesion protein that binds to the integrin receptors, is important for mechanotransduction (Atherton *et al*, 2016). It has have shown that the amino acid alanine at position 50 in vinculin N-terminal domain is critical for binding talin and that the alanine to isoleucine substitution (Vcl^A50I^) abolishes this interaction (Bakolitsa *et al*., 2004; Cohen *et al*., 2006). We generated GFP-tagged full length mutant (GFP-Vcl-FL^A50I^), GFP-Vcl-851^A50I^ and Vcl-258^A50I^ mutants and expressed it in cultured neocortical neurons. Interestingly, we found that the GFP-Vcl-FL^A50I^ did not affect axon growth when compared to Vcl-FL and empty vector (Fig. 2E, F). Similarly, the GFP-Vcl-851^A50I^ and Vcl-258^A50I^ mutants did not exacerbate the attenuated axon growth phenotype caused by Vcl-851 and Vcl-258 mutants respectively (Fig. 2F). We next asked whether talin binding is necessary for cell migration. For this we compared cell migration at E17.5 following expression of Vcl-FL^A50I^ and control GFP. We did not observe any difference in cell migration when Vcl-FL^A50I^ was expressed (Fig. 2G, H). Together our observations suggested that either talin-binding to vinculin is dispensable for axon growth and neuronal migration or that there are as yet unknown talin-binding residues in N-terminus of vinculin. It is also likely that other actin-binding proteins such as α-actinin, which has been shown to bind vinculin and integrins compensates for lack of talin binding (Le *et al*, 2017; Roca-Cusachs *et al*, 2013).

### Vinculin tail domain induces excessive branching and enlarged cell soma

Since Vcl-T affected neuronal morphology and migration, we sought to characterize further the functions of Vcl-T. We first looked at neurite branching and using Sholl analysis, we quantified the number of primary neurites arising from the cell soma and the number of secondary branches from the primary neurites. Compared to control neurons, we found that Vcl-T increased both the number of primary and secondary neurites (Fig. 3A, B). Neocortical neurons in culture have been shown to exhibit a stereotypic pattern of growth (Craig & Banker, 1994). Following plating, first the neurons extend several filopodial and lamellipodial protrusions from the cell soma (stage 1) followed by extension of many neurites (stage 2). Next, one of these neurites grows rapidly to become the axon (stage 3) followed by dendritic growth (stage 4). We therefore asked how Vcl-T affected the early stages of neuronal growth. For this, primary neocortical neurons were dissociated, transfected with Vcl-T or control GFP and cultured *in vitro*. Cells were fixed at 24 h and 48 h and analysed for number of primary and secondary neurites, and the length of the longest neurite. At 24 h post-plating, the Vcl-T transfected neurons showed greater number of primary neurites that also showed more branches (secondary neurites) with no significant difference in the length of the longest neurite (Suppl. Fig. 4A, B). At 48 h post-plating, the number of primary and secondary neurites remained higher in Vcl-T transfected neurons compared to control neurons (Suppl. Fig. 4C, D). However, the length of the longest neurite was shorter in Vcl-T expressing neurons compared to control (Suppl. Fig. 4C, D). Since, a single longer neurite was not evident in Vcl-T expressing neurons, we wondered whether neuronal polarity was affected in these cells. We immunostained the cultures for the axon specific marker, Tau. We found that although shorter, the Vcl-T expressing neurons extended only one axon (Fig. 3A, B). Together, these findings suggested that although Vcl-T expression does not affect neuronal polarity, Vcl-T attenuated axon growth and increased neurite branching from an early stage of neurite growth.

**Figure 3.**
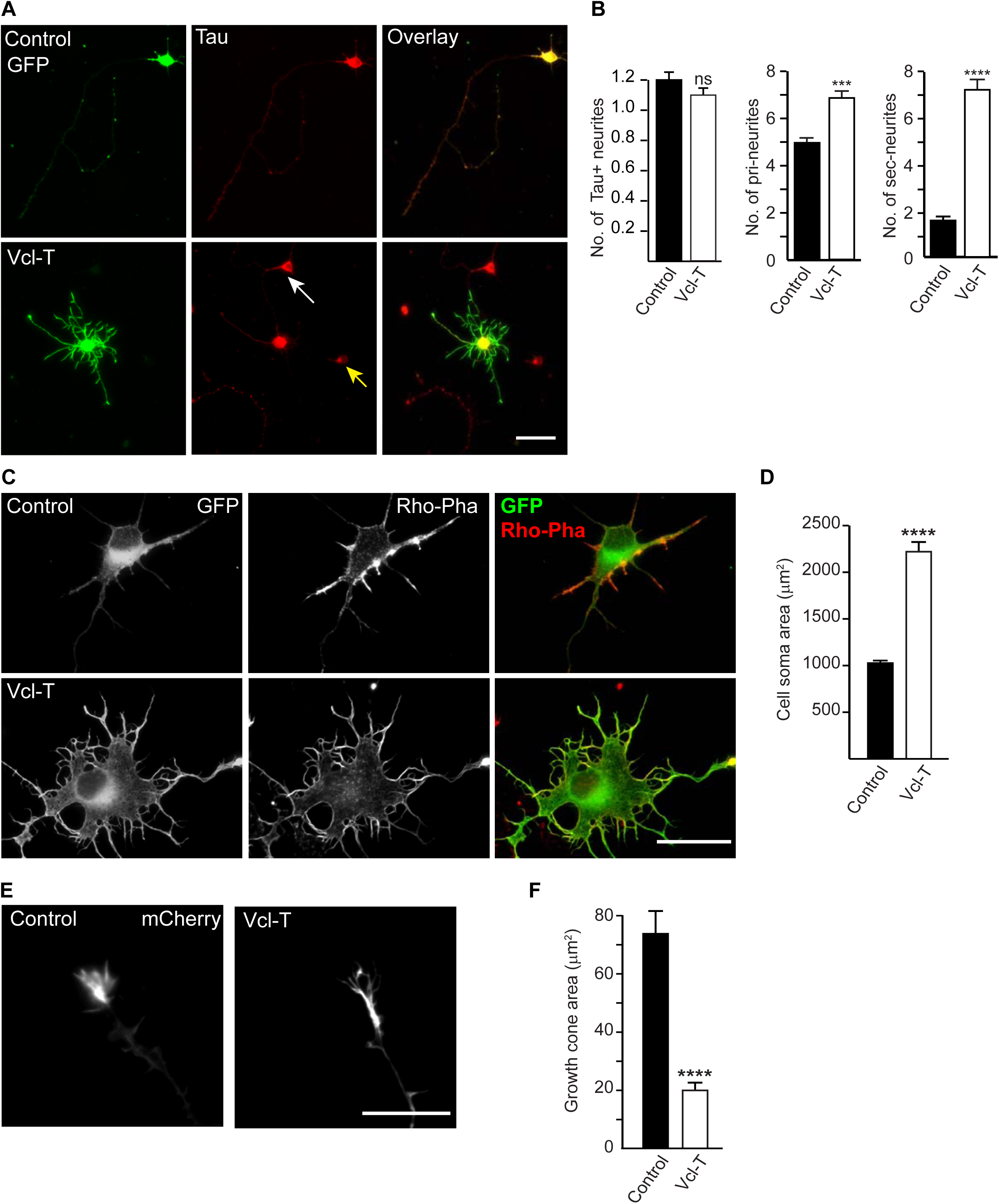
Vinculin tail domain increases neurite branching, cell soma area, and affects growth cone morphology *in vitro*. **(A)** Neocortical neurons from P0.5 pups were electroporated with Vcl-T and control GFP plasmid and grown for 4 DIV. Cells were fixed and immunostained for GFP (to label transfected neurons) and Tau (to label axons). Vcl-T expression increased both the number of primary and secondary neurites but did not affect polarity as seen by only one Tau^+^ neurite. Non-transfected neurons (white arrows) and dead cells (yellow arrows). **(B)** Quantification of number of Tau-positive neurites, and number of primary and secondary neurites from (A) (n=100 cells). **(C)** Representative images of neurons expressing either empty GFP (control) or Vcl-T plasmid at 4DIV. Neurons were immunostained for GFP and rhodamine-conjugated phalloidin (Rho-Pha) to visualize cell soma *in vitro*. Vcl-T expressing neurons had a wider spread of cell soma compared to control. **(D)** Quantification of the cell soma area in control and Vcl-T expressing neurons from (C) (n=100 cells). **(E)** Representative images from live imaging of control and Vcl-T expressing neurons co-transfected with LifeAct-mCherry to label actin. Vcl-T expression caused a smaller or collapsed growth cone morphology. **(F)** Quantification of the growth cone area in control and Vcl-T expressing neurons from (E) (n=25 cells). Scale, 50µm (A, C), 10µm (E). \*\*\**p*<0.001, \*\*\*\**p*<0.0001; ns, non-significant. Two-tailed student’s *t* test.

We had earlier observed abnormal cell soma morphology of Vcl-T expressing neurons *in vivo* (Fig. 2C, D). We therefore asked whether this phenotype is also seen in cultured neurons. Neocortical neurons were transfected with Vcl-T or control GFP and grown in culture for 4 days. Cells were fixed and stained for rhodamine-conjugated phalloidin to label F-actin. We found that Vcl-T expressing neurons exhibited a larger cell soma indicating that Vcl-T expression increased cell spreading (Fig. 3C, D). This is in contrast to observations made in cultured fibroblasts, wherein Vcl-T expression has been shown to reduce focal adhesions resulting in a smaller cell area (Humphries *et al*., 2007; Mierke *et al*, 2008).

Vinculin is present in the leading edge of processes where it helps in binding to the substratum to promote migration (Bays & DeMali, 2017). Since Vcl-T expressing neurons had shorter axons *in vitro* (Fig. 1A, B) and *in vivo* (Fig. 1C, D), we asked whether growth cone morphology is affected since improper growth cone can affect axon growth. For this, we analysed growth cone structure in the cultured neurons expressing Vcl-T and control GFP. We found that the area of the growth cone was smaller and appeared collapsed in Vcl-T compared to control GFP neurons (Fig. 3E, F). Together, these observations suggest that Vcl-T affects neuronal soma and growth cone morphology both *in vitro* and *in vivo*.

### Vcl-T-mediated branching phenotype requires Arp2/3 activity and actin-binding

The attenuation of axon growth and the excessive branching phenotype caused by Vcl-T suggested that vinculin tail domain inhibits F-actin polymerization (Le Clainche *et al*, 2010) and this in turn stabilizes the F-actin filament, thereby allowing the binding of other proteins to induce branching. The actin cytoskeleton binds to a large array of actin-binding proteins, which regulate the structural changes in actin including polymerization and nucleation. The actin-related protein 2/3 (Arp2/3) complex is an important actin nucleator that binds to existing actin filaments and causes branching of actin filaments and neurites (Blanchoin *et al*, 2000; Mullins *et al*, 1998; Sturner *et al*, 2019). We hypothesized that Vcl-T-induced neurite branching is mediated by the Arp2/3 complex. For this, we transfected neocortical neurons with Vcl-T or control GFP and cultured them in the presence of 25 nM of CK-666, a small molecule inhibitor of the Arp2/3 complex (Hetrick *et al*, 2013). Inhibition of Arp2/3 complex completely suppressed the branching phenotype and the number of primary and secondary neurites in Vcl-T expressing neurons was similar to that of control cells (Fig. 4A, B). However, inhibition of Arp2/3 complex failed to rescue the attenuated axon length caused by Vcl-T and caused further decrease in growth compared to Vcl-T expressing neurons (Fig. 4B).

**Figure 4.**
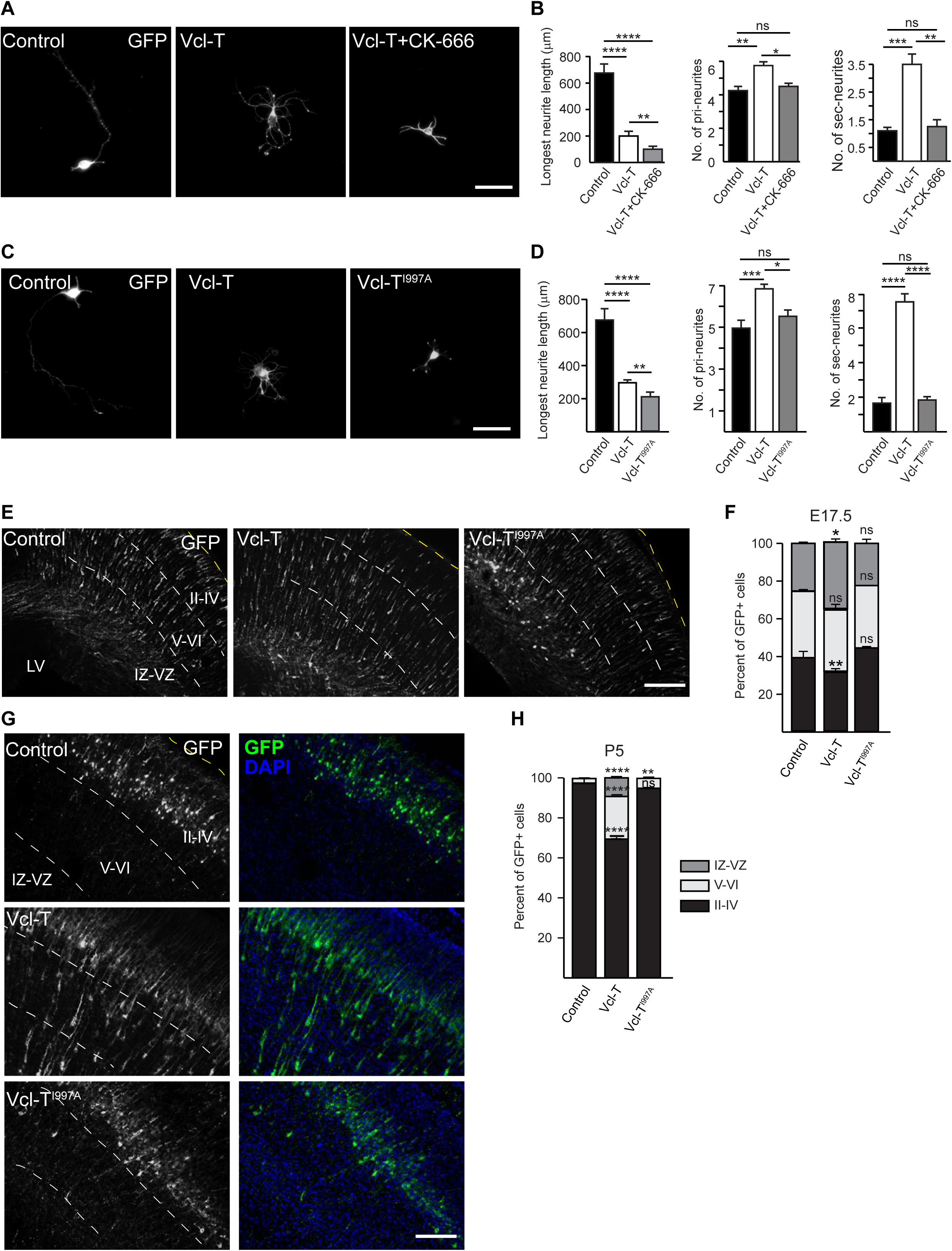
Blocking Arp2/3 complex and abolishing actin-Vcl-T interaction decreases branching but does not rescue axon length. **(A)** Representative images of neurons transfected with Vcl-T and treated either with DMSO (vehicle) or with 25nM CK-666 (Arp2/3 complex inhibitor) dissolved in DMSO. Vcl-T neurons exhibited enhanced neurite branching and this abolished in the presence of CK-666. **(B)** Quantification of longest neurite length, number of primary and secondary neurites of neurons from (A) (n=100 cells) **(C)** Neurons transfected with control, Vcl-T and Vcl-T defective in actin-binding (Vcl-T^I997A^) constructs and immunostained for GFP after 4DIV. Abolishing interaction of Vcl-T with actin reduced neurite branching to that seen in control cells. **(D)** Quantification of longest neurite length, number of primary and secondary neurites of neurons from (C) (n=300 cells). **(E)** Representative images of brain sections from E17.5 embryos following electroporation at E14.5 with control, Vcl-T and Vcl-T^I997A^ constructs. Abolishing actin-Vcl-T interaction rescues neuronal migration defect caused by Vcl-T. **(F)** Quantification of the percentage of GFP^+^ neurons in different layers of cortex from (E) (n=3 animals). **(G)** Analysis at post-natal day 5 (P5) also showed almost complete rescue of migration when actin-Vcl-T interaction is abolished. **(H)** Quantification of the percentage of GFP^+^ neurons in different layers of cortex and in IZ/VZ from (G) (n=3 animals). Scale, 50µm (A, C), 400µm (E, G). \*\**p*<0.01, \*\*\**p*<0.001, \*\*\*\**p*<0.0001; ns, non-significant. Two-tailed student’s *t* test in (B, D). One-way ANOVA and Tukey’s *post-hoc* test in (F, H).

The vinculin tail domain binds F-actin and crosslinks it into actin bundles (Huttelmaier *et al*, 1997; Menkel *et al*., 1994). Among several amino acid residues identified in Vcl-T domain that bind to actin, the isoleucine 997 to alanine mutant (Vcl-T^I997A^) was found to be the most impaired in actin binding (Thompson *et al*, 2014). We generated Vcl-T^I997A^ to determine the effect of abolishing actin-Vcl-T interaction on axon growth and neuronal morphology. Neocortical neurons expressing Vcl-T^I997A^ showed a decreased branching phenotype compared to Vcl-T expressing neurons and the number of primary and secondary neurites were comparable to that of control neurons (Fig. 4C, D). But similar to that seen with Arp2/3 inhibition, abolishing Vcl-T-actin interaction did not rescue axon growth and in fact, it exacerbated the stunted axon phenotype (Fig. 4D).

We next asked whether abolishing Vcl-T-actin interaction can rescue the enlarged cell soma phenotype. We found that Vcl-T^I997A^ mutant did not cause any increase in cell soma area as seen in Vcl-T expressing neurons (Suppl. Fig. 5A, B). This suggested that Vcl-T expression affected actin dynamics resulting in an enlarged soma and abolishing this interaction with actin prevented enlargement of cell soma. We had earlier shown that Vcl-T caused a reduction in the area of the growth cone and this could be due to ability of Vcl-T to bind the barbed ends of F-actin preventing its polymerization (Fig. 3E, F) (Le Clainche *et al*., 2010). We therefore asked whether abolishing Vcl-T-actin binding can rescue the growth cone morphology and promote axon extension. For this, we transfected neurons with control GFP plasmid, Vcl-T or Vcl-T^I997A^ along with LifeAct-mCherry to label actin. To assess growth morphology, Vcl-T^I997A^ expressing neurons were imaged for LifeAct-mCherry expression. We found that the growth cone morphology was not rescued in the absence of actin binding suggesting that either Vcl-T binding recruits other actin-binding proteins that inhibit axon extension or that the head and neck domains are required for normal axon growth (Suppl. Fig. 5C, D).

Since abolishing actin-binding in Vcl-T (Vcl-T^I997A^) was able to decrease Vcl-T-induced branching, we were interested in knowing how the full-length vinculin that is deficient in actin binding would function in neurons. For this we generated a similar isoleucine 997 to alanine point mutation in full-length vinculin (GFP-Vcl-FL^I997A^) and expressed it in neurons. We found that Vcl-FL^I997A^ expression affected axon growth to a moderate but a significant extent. However, it did not induce or affect any branching (Suppl. Fig. 6).

### Abolishing actin binding rescues neuronal migration defects caused by Vcl-T

We had seen that Vcl-T impaired neuronal migration *in vivo* (Fig. 1E, F). We next sought to determine whether abolishing actin interaction will rescue this defect in migration. For this, we electroporated Vcl-T, Vcl-T^I997A^, control plasmid into E14.5 embryos and visualized neuronal migration three days later at E17.5. In contrast to the slower migration rate exhibited by Vcl-T expressing neurons, the Vcl-T^I997A^ neurons showed normal migration and were comparable to that seen in control embryos (Fig. 4E, F). We next looked at neuronal migration at a later time point (P5) following electroporation at E14.5. In control GFP transfection, the neurons had migration to the upper layers as expected while the Vcl-T expressing neurons were stalled in the lower layers (Fig. 4G, H). However, the Vcl-T^I997A^ expressing neurons exhibited normal migration to the upper layers (Fig. 4G, H) indicating that abolishing actin binding rescued migration defect caused by Vcl-T. This was further confirmed by immunostaining for the upper layer marker, Cux1. In control GFP and Vcl-T^I997A^ transfected embryos, all Cux1^+^ cells were seen in layers 2 to 4 suggesting normal migration. However, in Vcl-T transfected embryos, many Cux1^+^ neurons were seen in the lower layers indicating retarded migration (Suppl. Fig. 7). Vcl-T expression was seen to impair axon growth *in vivo* (Fig. 1C, D). We asked whether Vcl-T^I997A^ mutant can rescue the axon growth defect caused by Vcl-T. For this, we analysed the length of the axon along the corpus callosum at P5. In control GFP transfected brains, the axons from the transfected hemisphere had crossed over to the contralateral hemisphere while in the Vcl-T transfected embryos, the axons were shorter and very few crossed the midline (Fig. 5A, B). However, in Vcl-T^I997A^ transfected brains, the axons were longer and crossed the midline but were significantly shorter compared to control neurons (Fig. 5A, B). Together, these observations suggested that abolishing actin binding was able to completely rescue the migration defect caused by Vcl-T but could only moderately rescue axon growth *in vivo*.

**Figure 5.**
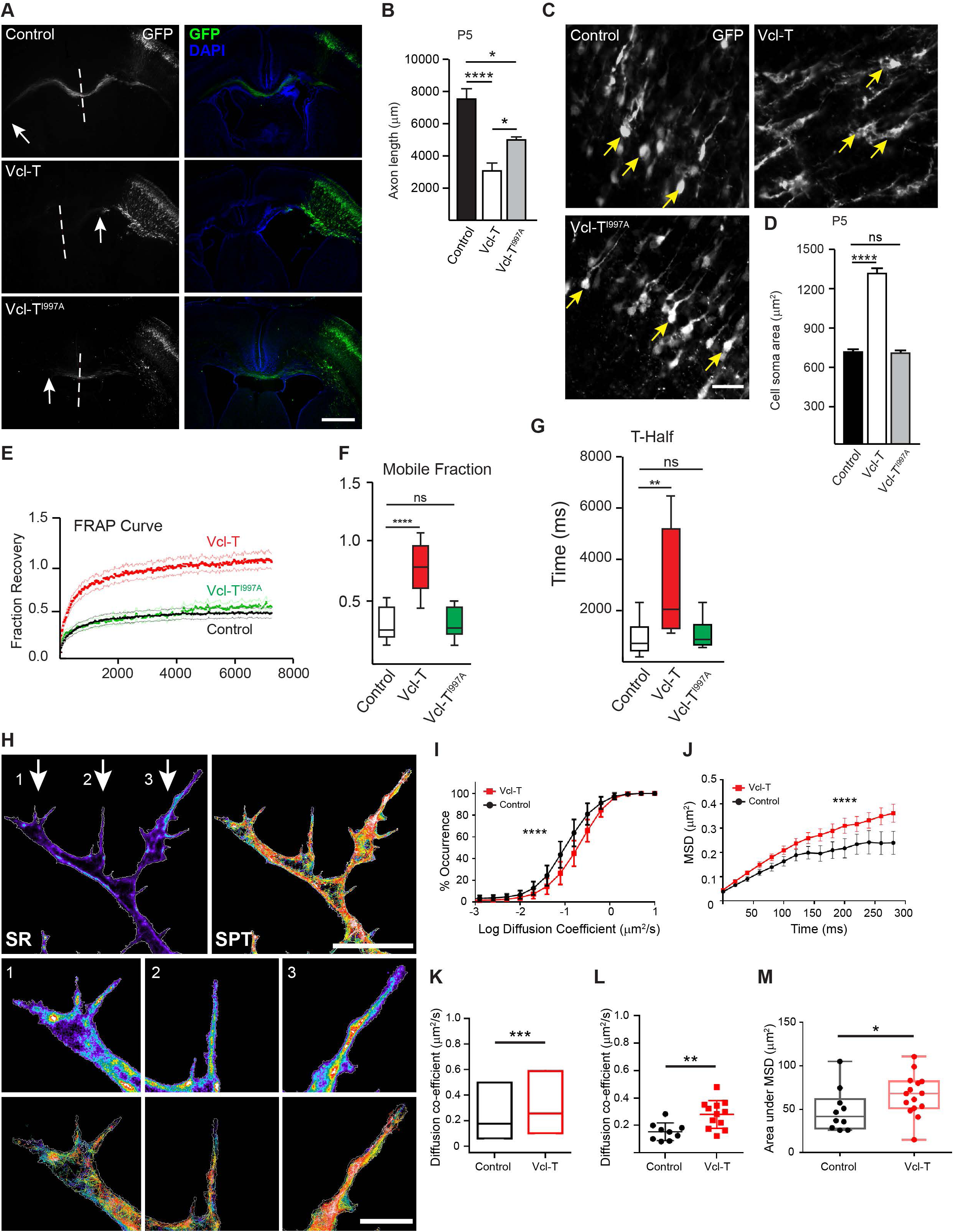
Abolishing actin-Vcl-T interaction reduces cell soma area *in vivo* and Vcl-T increases actin mobility *in vitro*. **(A)** Representative images of brain sections electroporated with control, Vcl-T and Vcl-T^I997A^ constructs. Vcl-T^I997A^ defective in actin-binding only partially rescues axon growth deficit caused by Vcl-T. Arrows indicate the axon terminals along the corpus callosum. Dashed lines represent the longitudinal fissure separating the two cerebral hemispheres. Scale, 2mm. **(B)** Quantification of length of axons from (A) (n=3 animals. \**p*=0.0193, Vcl-T-Vcl-T^I997A^; \**p*=0.0170, Ctrl-Vcl-T^I997A^. **(C)** Representative images of brain sections showing cell soma morphology of neurons expressing control GFP, Vcl-T and Vcl-T^I997A^. Vcl-T expressing neurons exhibited abnormal soma (arrows) compared to control neurons, and this was rescued when actin-Vcl-T interaction was abolished. Scale, 150µm. **(D)** Quantification of neuronal cell soma area from (C) (n=100 cells, n=3 animals). \*\*\*\**p*<0.0001; ns, non-significant. **(E-G)** Neurons were transfected with control GFP, Vcl-T or Vcl-T^I997A^ along with LifeAct-mCherry to visualize actin. Fluorescence recovery curve **(E)**, mobile fractions **(F)** and half-time **(G)** of recovery of fluorescence of LifeAct-mCherry after FRAP of neurons transfected with control, Vcl-T and Vcl-T^I997A^ constructs, showing that neurons expressing Vcl-T have a higher mobile fraction and higher t-half of actin compared to control neurons. Abolishing actin-Vcl-T interaction (Vcl-T^I997A^) reduces the mobile fraction and reduces t-half to normal level. (n=10 cells). \*\**p*<0.01, \*\*\*\**p*<0.0001; ns, non-significant. **(H-M)** Single particle tracking of mEOS3.2-LifeAct in control and Vcl-T expressing neurons. **(H)** Representative super-resolution intensity map reconstructed from single molecule localization (left) and single particle trajectories (right) generated from neurons expressing mEOS3.2-LifeAct. White arrows indicate the extensions of the neurite. Scale, 10 µm. The regions marked by arrows are represented below. For each region indicated by arrow, the magnified super-resolution intensity map (SR) and single particle trajectories (SPT) for the same region is shown. Scale, 2 µm. (**I**) Normalized cumulative distribution of logarithmic values of instantaneous diffusion coefficient of the generated tracks (Mean +/-SEM). \*\*\*\**p*<0.0001 (Kolmogorv-Smirnov test). (**J**) Plot of average mean squared displacement (MSD) of all trajectories above 6 frames for control and Vcl-T. \*\*\*\**p*<0.0001 (Wilcoxon matched-pairs signed rank test). (**K**) Pooled values of instantaneous diffusion coefficient of all the detected trajectories from control and Vcl-T (median, IQR). \*\*\**p*=0.0002 (Unpaired t test). (**L**) Distribution of median diffusion coefficients of each neuron compared across the conditions. \*\**p*=0.0039 (Unpaired t test). (**M**) Distribution of area under curve for average MSD for trajectories obtained from each cell of control versus Vcl-T. \**p*=0.0469 (Mann-Whitney test).

Vcl-T expression also resulted in an enlarged cell soma *in vivo* (Fig. 2C, D). We next asked whether inhibiting Vcl-T-actin binding can rescue the enlarged cell soma phenotype. For this, we measured the cell body area of GFP^+^ neurons in the upper layers of the cortex at P5 in brain sections transfected with Vcl-T^I997A^ and compared the results with that of Vcl-T and control GFP expressing neurons. We found that the cell soma area of Vcl-T^I997A^ expressing neurons was comparable to that of control neurons and this was similar to that observed in *in vitro* culture experiments (Fig. 5C, D). Together these results suggested that the defects in migration and abnormal morphology of neurons caused by vinculin tail domain was due to abnormal interaction with actin and that abolishing this interaction could rescue both these defects *in vivo* and *in vitro* but not axon length.

### G-actin mobility is faster in neurons expressing vinculin tail domain

To understand the molecular mechanisms underlying the increased branching phenotype caused by Vcl-T, we studied actin mobility in cultured neurons. The Vcl-T^I997A^ expression data indicated that actin-binding is necessary for the branching phenotype. Since Vcl-T has been shown to bind the barbed ends of F-actin and prevent its polymerization (Le Clainche *et al*., 2010), we hypothesized that this would increase the G-actin pool, thus making Arp2/3 complex to nucleate and generate branching. To test this hypothesis and to visualize actin in real-time, we co-electroporated neurons with LifeAct-mCherry along with control GFP or Vcl-T construct. LifeAct, the first 17 amino acids of actin-binding protein 140 from *S. cerevisiae*, binds to actin filaments without modifying its function (Riedl *et al*, 2008). mCherry was tagged to LifeAct for visualization of actin in cells. To study the behaviour of actin in neurons, we photobleached mCherry in a small region close to the edge of the longest neurite by using a high-power laser of 561 nm wavelength and monitored fluorescence recovery over time (FRAP). The fluorescence recovery was fit as a single exponential function of time and half-time of recovery and mobile fraction was calculated from the curve fitting (Fig. 5E-G). In Vcl-T expressing neurons there was recovery of a higher mobile fraction of LifeAct (0.7655 +/- 0.06361) when compared to mobile fraction of control cells (0.3071 +/- 0.03738) (Fig. 5F). We also observed a significant increase in t-half of recovery in presence of Vcl-T (3227ms +/- 647.2ms) compared to control (941.5ms +/- 171.0ms) (Fig. 5E). This increase in mobile fraction could be due to either an increase in G-actin pool or impaired F-actin dynamics (actin treadmilling). Interestingly, neurons expressing the Vcl-T^I997A^ mutant, that is defective in actin binding, showed a complete rescue in the mobile fraction (0.3268 +/- 0.03837) and t-half (1139ms +/- 206.2ms) of actin (Fig. 5F, G).

To further characterize the effect of Vcl-T on actin mobility, we expressed Vcl-T along with mEOS3.2-LifeAct (Zhang *et al*, 2012). Using photoactivation properties of mEOS3.2, we performed single particle tracking based photoactivation localization microscopy on LifeAct in transfected cells. High-resolution intensity map and single particle trajectories were generated for each neuron (Fig. 5H). Instantaneous diffusion coefficient (D) and mean squared displacement (MSD) were calculated and used to estimate diffusion dynamics and confinement of the LifeAct molecules. The trajectories displayed a wide range of diffusion coefficient and a significant difference in distribution of logarithmic values of diffusion coefficient was observed, indicating that mEOS3.2-LifeAct was more mobile in presence of Vcl-T (Fig. 5I). In presence of Vcl-T, instantaneous mEOS3.2-LifeAct showed increase in diffusion coefficient (0.2570 µm2/sec, IQR: 0.1012 - 0.5867 µm2/sec) when compared to control (0.1749 µm2/sec, IQR: 0.06114 - 0.4980 µm2/sec) after pooling all the trajectories obtained (Fig. 5K). To rule out the possibility of cell to cell variation in diffusion coefficient, we compared the median of diffusion coefficient from all cells extracted individually and found the difference to be significant (Fig. 5L). These values were consistent with the pooled analysis. The average MSD of LifeAct saturated with time in both conditions indicating confined diffusion in both cases. However, the saturation values of LifeAct in the presence of Vcl-T were higher than the control indicating lesser confinement compared to control (Fig. 5J). This was also validated by significant difference in the area under the curve for MSD, for the average MSD curves generated for each cell (Fig. 5M). Together these observations confirm that the actin molecules became less confined and more mobile in the presence of vinculin tail domain. This could facilitate nucleation of actin and branching by the Arp2/3 complex resulting in enhanced branching induced by the Vcl tail domain.

Here, we demonstrate that vinculin plays a critical role in neuronal migration *in vivo* and in axon growth *in vitro* and *in vivo*. Although full length vinculin is required for optimal axon growth, the mutants used in this study provide novel insights into some of the functional domains of vinculin. The constitutively active vinculin greatly enhanced axon growth both *in vitro* and *in vivo*. This observation opens up the question of whether vinculin activation can promote axon growth on inhibitory substrates and also raises the possibility that inhibitory substrates could likely affect axon growth by blocking vinculin-dependent actin dynamics. We also observed that preventing vinculin-talin binding did not affect migration and axon growth indicating that this interaction is dispensable for these vinculin-dependent functions, in contrast to that seen in fibroblasts. Interestingly, vinculin tail domain affected neuronal morphology in several ways compared to the other functional domain mutants of vinculin. Among these was a striking increase in neurite branching and enlarged cell soma, with the latter phenotype being different than that seen with fibroblasts, in which Vcl-T expression resulted in a smaller cell. These phenotypes could be rescued by abolishing interaction of the tail domain with actin and by inhibiting the Arp2/3 complex. Vcl-T, by binding to the barbed end of F-actin prevents polymerization and thus resulting in an increase in mobile actin pool. This in turn stabilizes F-actin and causes an Arp2/3-dependent branching and the process continues, resulting in highly branched but shorter neurites. It will be interesting to see how these Vcl-T will affect the localization and properties of the other actin-binding proteins that play a role in axon growth.

## Materials and methods

### Animals

For all experiments, C57Bl/6J mice were used. All experiments were conducted in accordance with the animal care standards of the Institutional Animal Ethics Committee of the authors’ institution.

### Constructs

Sequences of guide RNAs (gRNAs) for deleting Vcl gene was selected using DeskGen software (www.deskgen.com) and two gRNAs that bind to flanking regions of translational start site were chosen. The chosen gRNA sequences are as follows: Vcl-gRNA-1, 5’-GCGTACGATCGAGAGCATCC-3’ and Vcl-gRNA-2, 5’-GCTAGCGGGGCGGCGTACCG-3’. The gRNAs were first individually cloned into pX330-U6-BB-CBh-hSpCas9 plasmid (Addgene No: 42230) using BbsI site. These two gRNAs were then subcloned into pSpCas9n(BB)-2A-GFP plasmid (pX461; Addgene No: 48140). We made use of Cas9 nickase (Cas9n) to decrease any potential OFF-target effects. The resultant vector was named pX461-Vcl-sg1+2--Cas9n-2A-GFP. pSpCas9n(BB)-2A-GFP (PX461) and pX330-U6-Chimeric_BB-CBh-hSpCas9 were a gift from Feng Zhang (Addgene plasmid # 48140, 42230; http://n2t.net/addgene:48140; RRIDs: Addgene_48140, Addgene_42230).

The mouse *Vcl*-encoding gene (Vcl-FL) was amplified from cDNA synthesized from whole brain mRNA. Vcl-FL^A50I^ was generated from Vcl-FL by mutating the alanine residue at 50th position to isoleucine using site-directed mutagenesis (SDM). Vcl-FL^I997A^ was generated from Vcl-FL by mutating isoleucine residue at 997th position to alanine using SDM. Vcl-LD was generated from Vcl-FL by mutating 1060th residue from arginine to glutamine and residue at 1061st from lysine to glutamine by using SDM. Vcl-258 was amplified from full length vinculin by using the following primers: Forward 5’-CTTGAATTCGATGCCGGTGTTTCACACG-3’ and Reverse 5’-CGGGGTACCCCAGGCGTCTTCATCCC-3’. Vcl-258^A50I^ was generated from Vcl-258 by mutating alanine residue at 50th position to isoleucine using SDM. Vcl-851 was amplified from full length vinculin using following primer: Forward, 5’-CTTGAATTCGATGCCGGTGTTTCACACG-3’ and Reverse, 5’-CGCGGTACCCTCTTCTGGTGGTGGGG-3’. Vcl-851^A50I^ was generated from Vcl-851 by mutating alanine residue at 50th position to isoleucine using SDM. Vcl-T12 was generated by mutating residues 974th from glutamate to alanine, 975th from lysine to alanine, 976th from arginine to alanine and 978th from arginine to alanine from full length vinculin by using SDM. Vcl-T was amplified from Vcl-FL by using these primers: Forward 5’-CTTGAATTCGAAGGATGAAGAGTTCCCTG-3’ and Reverse 5’-CGCGGATCCCTGGTACCAGGGAGTCTTTC-3’. All these clones were generated in pCAG-C1 vector. pCAG-C1 was generated from pEGFP-C1 vector by replacing the CMV promoter by CAG promoter from pCAGEN, a gift from Connie Cepko (Addgene plasmid # 11160; http://n2t.net/addgene:11160; RRID: Addgene_11160). Vcl-T^I997A^ clone was generated from Vcl-T by site-directed mutagenesis of the 997th amino acid residue from isoleucine to alanine. LifeAct-mCherry was a gift from Dr. Paul Bridgman, Washington University School of Medicine, St. Louis, MO. mEos3.2-C1 was a gift from Michael Davidson & Tao Xu (Addgene plasmid # 54550; http://n2t.net/addgene:54550; RRID: Addgene_54550). Vcl-T-mEOS3.2-LifeAct was generated by sub-cloning a Vcl-T-T2A fragment in mEOS3.2-LifeAct clone.

### Site-directed mutagenesis

Complementary primers of 30-40 bp were designed with the mutation site in the middle of the sequence and 50 ng of template DNA was used along with 25 pmoles each of the mutant primers. 2 min extension time for every 1 kb of template length was used. PCR was for a total of 18 cycles and the following cycling conditions were used: 95°C for 3 min; 68°C for (2 min/kb); 18 cycles; 72°C for 10 min; end at 4°C. The reaction mixture was digested with DpnI enzyme to remove template DNA and ligation was set with T4DNA ligase for 1 h at room temperature. A small aliquot of the ligation reaction was transformed into DH10B competent cells and transformants were selected on LB+antibiotic plates. The site-directed mutation was confirmed by DNA sequencing of both strands of at least 2 independent clones.

### Immunocytochemistry

Neurons were fixed using 4% PFA + 4% sucrose in PBS for 20 min and washed in PBS, 2×10 min. The cells were permeabilized with 0.1% Triton-X100 in PBS for 5 min and blocked with 10% BSA in PBS for 30 min at room temperature with gentle rocking. Primary antibody incubation was done in 3% BSA for 1 hour at 4 °C with gentle rocking and then washed with PBS, 3×2 min. Secondary antibody incubation was done in 3% BSA for 30 min at room temperature, washed with PBS, 3×2min and mounted with VECTASHIELD mounting medium with DAPI (#H-1000, Vector Laboratories, USA). The following primary antibodies were used: anti-GFP (#GFP-1020, 1:1000, Aves labs, Inc. USA), anti-β-III tubulin (#Tuj, 1:1000, Aves labs, Inc. USA), anti-Tau (#314002, 1:1000, Synaptic Systems, Germany). The following secondary antibodies from Invitrogen Inc., were used: AlexaFluor (AF)-488 conjugated anti-chicken (# A11039, 1:1000), AF-594 conjugated anti-mouse (#A21203, 1:1000), AF-594 conjugated anti-rabbit (# A11012, 1:1000).

### Phalloidin staining

Rhodamine-phalloidin (#6876, 1:125, Setareh Biotech, USA) was added along with secondary antibody during immunocytochemistry.

### Immunohistochemistry

Immunohistochemistry was done as follows: embryonic or post-natal brains were fixed by transcardial perfusion using ice-cold PBS followed by 4%PFA, post-fixed in 4% PFA overnight and then cryopreserved in 30% sucrose until the brains were saturated with sucrose and sunk to the bottom of the tube, and frozen at -80° C until use. The frozen brains were sectioned at 20 μm thickness, collected on Superfrost plus slides (Brain Research Laboratories, USA) and dried overnight before staining. Blocking/permeabilization was done in blocking solution (3% bovine serum albumin + 0.3% Triton-X100 + 1% goat serum in PBS) for 1 h at RT. Primary antibody incubation was done in the blocking solution overnight at 4 °C with gentle rocking and washed with PBS for 5×5 min. Secondary antibody was incubated in the blocking solution for 1 h at RT, washed with PBS for 5×5 min and slides were mounted with VECTASHIELD mounting medium with DAPI (Vector Laboratories, USA). The following primary antibodies were used: anti-GFP (GFP-1020, 1:1000, Aves labs, Inc. USA), anti-Cux1 (#sc-13024, 1:50, Santa Cruz Biotechnology, USA). The following secondary antibodies from were used: AlexaFluor-488 conjugated anti-chicken (#A11039, 1:1000,), AlexaFluor-594 conjugated anti-rabbit (#A11012, 1:1000).

### Neuronal culture and transfection

Neocortex from C57Bl/6 pups (postnatal day 1-3) was dissected in Hibernate-A medium (#A1247501, Gibco, USA), digested with papain (#LS003124, Worthington, USA) for 30 min at 30° C and dissociated with micropipette tips in Neurobasal-A medium (#10888022) supplemented with glutaMAX (#35050061, Gibco, USA), B27 (#17504044, Gibco, USA) and 33 U/ml penicillin/streptomycin. The tissue was gently titrated, cells were pelleted by centrifugation, and washed twice with Hibernate-A medium. Cells were then plated at a density of 125 cells/mm^2^ on coverglasses (#41001112, Glaswarenfabrik Karl Hecht, Germany) coated with poly-L-lysine (# P2636, Sigma, USA) and laminin (#354232, BD, USA). Poly-L-lysine (50 µg/ml) coating was done for 2 h at 37 °C, washed thrice with Milli-Q water and dried at RT for 15 min followed by laminin (10 µg/ml) coating for 2 h at 37 °C. Cultures were incubated at 37°C with 5% CO2. After allowing for cells to attach to poly-L-lysine coverglass (20 min post-plating), the medium was gently exchanged with new neuronal culturing medium.

After 4 days in culture, neurons were fixed in 4% PFA+4% sucrose in PBS for 20 min. Neuronal transfection was done just before plating using Nucleofection (Lonza, Switzerland) according to the manufacturer’s instructions.

### Drug treatment

For inhibiting Arp2/3 protein complex *in vitro*, the pharmacological inhibitor CK-666 (Sigma Aldrich, SML0006) was used at a concentration of 25nM. CK-666 or DMSO (control) was added at 24 h after plating the neurons and neurons were allowed to grow for more 3 days.

### In utero electroporation

*In utero* electroporation for different mutants constructs of *Vcl* were done on embryonic day 14.5 (E14.5) in C57bl/6J embryos by injecting 1-2μl of 1μg/μl DNA in the lateral ventricle of one of the cerebral hemispheres. Electroporation was done using ECM 830 square wave electroporation system (BTX, USA) using 5 mm tweezertrode. The following parameters were used: voltage, 50V; pulse length, 50 milliseconds; number of pulses, 5 pulses; pulse interval, 100 milliseconds. Brains were harvested at either embryonic day 17.5 (E17.5) or at post-natal day 5 (P5) for immunostaining.

### Fluorescence Recovery After Photobleaching (FRAP)

FRAP was performed on an inverted fluorescence motorized fluorescence microscope (IX83 Olympus, Japan) equipped with a 100X 1.49 NA PL-APO objective and maintained at 37°C with the aid of a whole microscope temperature-controlled chamber. Neuronal primary cultures were loaded in open chamber (Ludin chamber, Life Imaging Services, Switzerland). The image acquisition was controlled by Metamorph (Molecular devices). Image acquisition was performed by illuminating with 488 or 561 nm laser and fluorescence were collected with filters to observes Green and Red fluorescence proteins in total internal reflection microscopy mode at 50Hz (Gataca Systems, France). Photo bleaching was done by using 561nm laser line for 60 ms in a circular region of diameter 20 pixels close to the end of the longest neurite. For baseline normalization, 10 images were acquired before photobleaching. To observe the recovery of fluorescence 400 frames were acquired after photobleaching sequence. For comparison between different experiments average of prebleached values were normalized to zero. Percentage calculation and normalization were performed using Microsoft Excel software. The recovery of fluorescence was fit to One-Phase association equation in GraphPad Prism. The quantification of recovery of fluorescence, mobile fraction and T-half graphs were generated from the curve fitting.

### Single Particle Tracking (SPT)

Single particle tracking using photoactivation was performed on live neurons expressing mEOS3.2 as previously described (Nair et al., 2013). These cells were loaded in an open chamber and photoactivated using a 405nm laser (Omicron, Gataca Systems, France) and the images were acquired using 561nm laser (Cobolt, Gataca Systems, France). The power of lasers was adjusted to keep the number of the stochastically activated molecules constant and well separated during the acquisition. Continuous acquisition of 20000 frames of images at 50Hz (20ms) were captured using a sensitive EMCCD camera (Evolve, Photometric). Image acquisition was controlled using MetaMorph software (Molecular Devices). Super resolved intensity images and single particle trajectories were obtained using custom built analysis for high density single particle tracking which were described previously (Nair *et al*, 2013).

### Data analyses and quantifications

Western blot quantification was done by using ImageJ. Longest neurite length was measured using the ‘traced line’ tool in MetaMorph software (Molecular Devices).

Numbers of primary and secondary neurites in cultured neurons were counted using Sholl analysis for different *vinculin* mutants. Cell body area was measured using ‘trace area’ tool in MetaMorph software (Molecular Devices). Numbers of cells in different layers of cortex was counted in ImageJ by using ‘cell counter’ plugin. All the graphs were made by using GraphPad Prism software.

### Statistical analyses

One-way ANOVA followed by Tukey’s *post hoc* test or two-tailed Student’s *t* test was used to determine statistical differences between groups using GraphPad Prism software. Error bars in figures are mean ± SEM unless otherwise stated. For statistical difference between frequency distribution of logarithmic values of diffusion coefficients we used Kolmogorov-Smirnov test. Diffusion coefficients were compared by unpaired *t* test. Wilcoxon matched-pair signed rank test was used to conclude the difference between MSD curves. Area under curve was calculated and compared using Mann-Whitney test.

### Image acquisition

Fluorescent images were acquired using Nikon Eclipse 80i upright fluorescent microscope (Nikon Instruments, USA), and confocal LSM880 microscope (Zeiss, Germany). Images were acquired and pseudo coloured using MetaMorph software (Molecular Devices) for Nikon Eclipse 80i microscope and Zen software for confocal LSM880 microscope.

## Supplementary Figure Legends

**Suppl. Figure 1.**
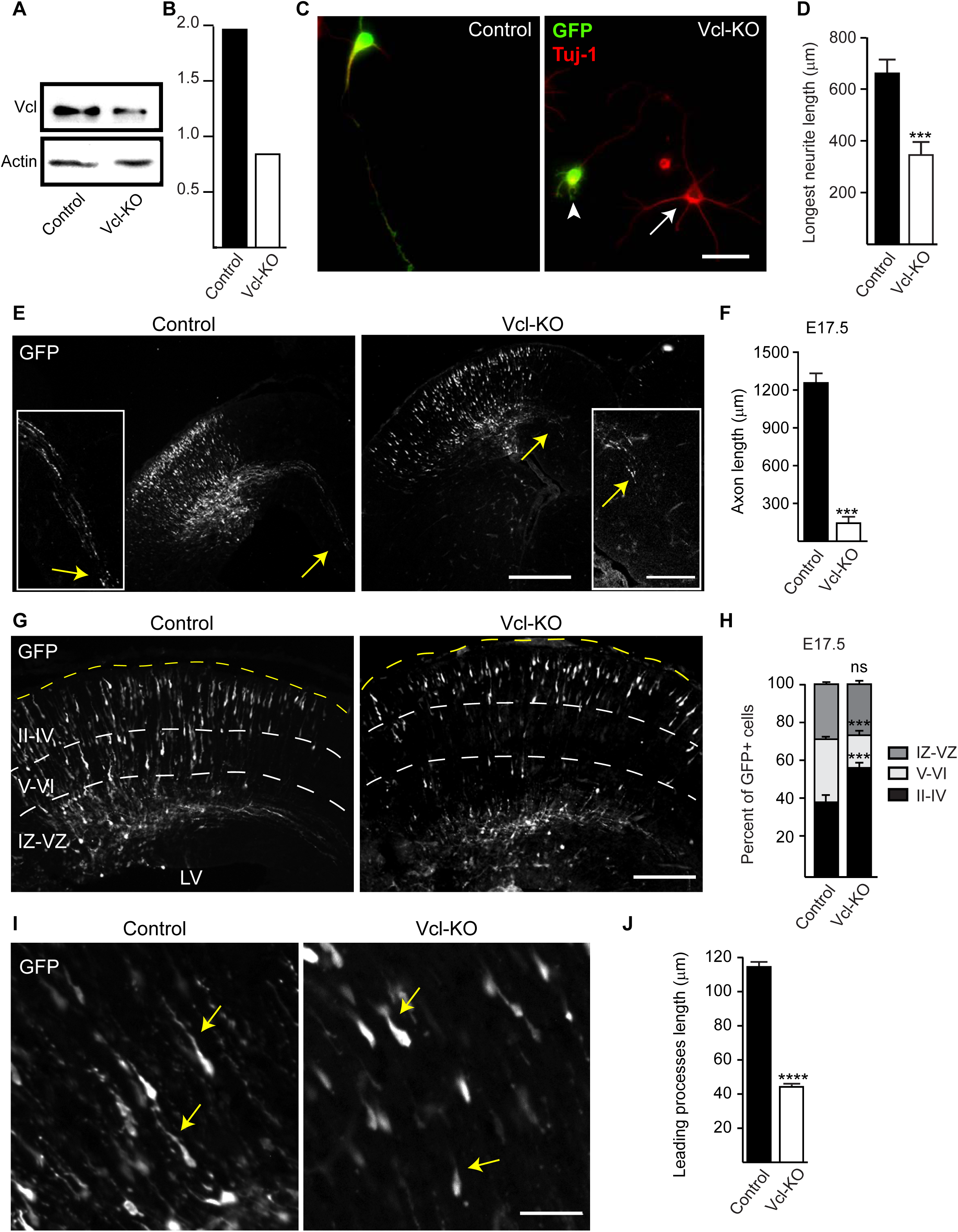
*Vinculin* deletion attenuated axon growth and enhanced neuronal migration. **(A)** NIH3T3 cells were transfected with a pair of single guide RNAs (gRNA) targeting *Vinculin* along with Cas9n nickase and cells were lysed at 72 hrs post-transfection. Western blot showing reduction in vinculin expression in Vcl-gRNA transfected cells (Vcl-KO). Actin was used for loading control. **(B)** Quantification of western blot signals from (A). **(C)** Primary neocortical neurons were electroporated with either empty Cas9n-T2A-GFP plasmid or Vcl-gRNA-Cas9n-T2A-GFP construct. Cells were fixed at 4 DIV and immunostained for GFP for transfected neurons and β-III-tubulin (Tuj-1) to visualize neurites. Neurons expressing Vcl-gRNAs exhibited shorter neurites compared to control neurons. Non-transfected (arrows) and Vcl-gRNA expressing cells (arrowhead) are indicated. **(D)** Quantification of longest neurite length from (C) (n=100 cells). **(E)** Neuronal progenitors at E14.5 were electroporated with either empty GFP (control) vector or Vcl-gRNA construct at E14.5 and analysed at E17.5 showed that Vcl-KO stunts axonal growth *in vivo* as well. Arrows indicate the axon terminals along corpus callosum. Inset shows magnified view of the axonal fibre. **(F)** Quantification of axon length from (E) (n=3 animals). **(G)** Embryos were electroporated at E14.5 with Vcl-gRNAs and analysed at E17.5. Vinculin deletion enhances neuronal migration during corticogenesis. **(H)** Quantification of the percentage of GFP^+^ neurons in different layers of cortex shown in (G) (n=3 animals). **(I)** Analysis at E17.5 also showed that vinculin deletion stunts leading processes of neurons. **(J)** Quantification of the length of leading processes from (I) (n=200 cells, n=3 animals). Scale 50µm (C), 2mm (E), 150 µm (inset in E), 400µm (G), 100µm in (I), \*\*\**p*<0.001, \*\*\*\**p*<0.0001. ns, non-significant. Two-tailed student’s *t* test (D, F, J). One-way ANOVA and Tukey’s *post-hoc* test (H).

**Suppl. Figure 2.**
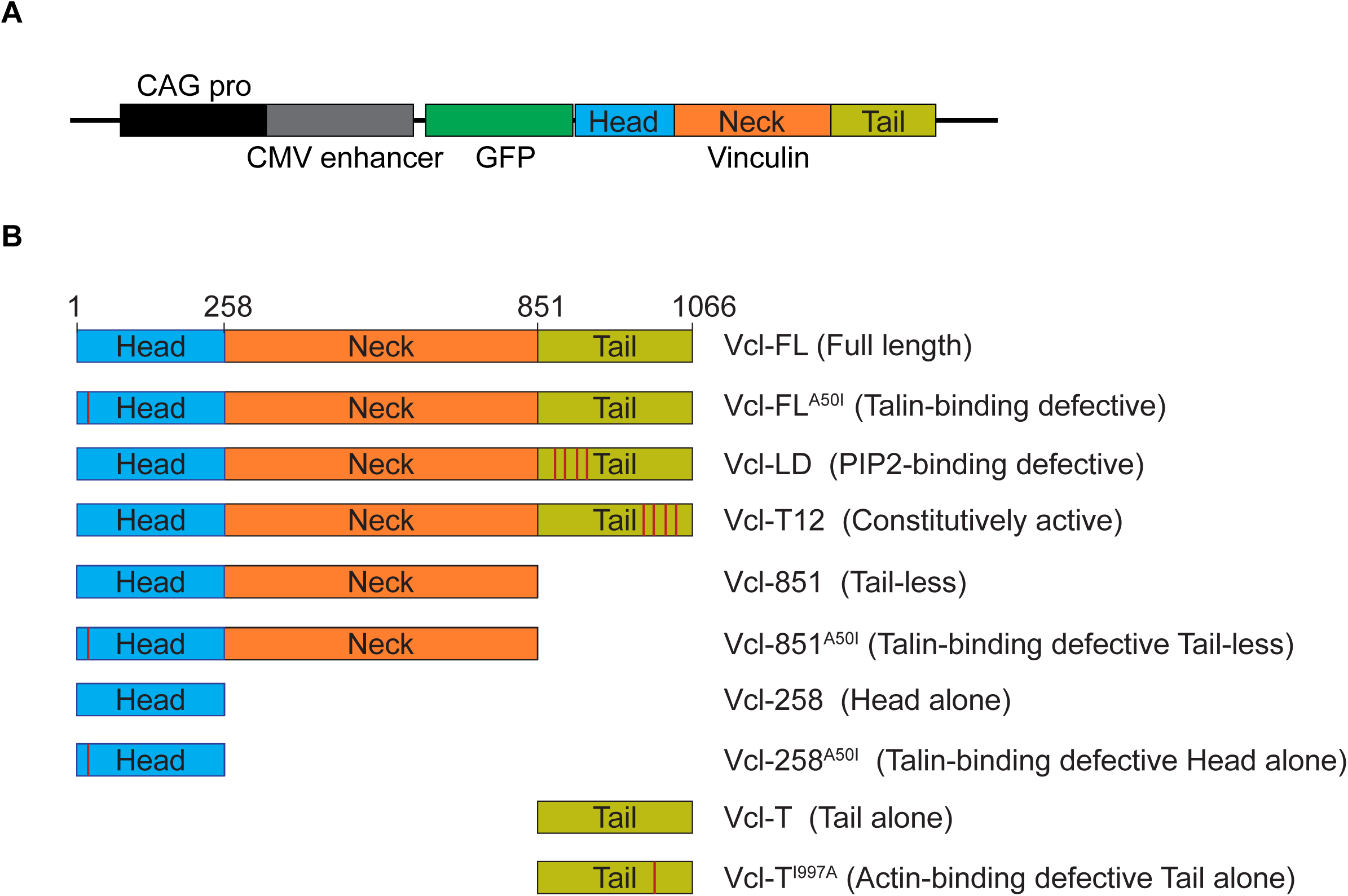
**(A)** Cartoon showing the partial vector map of full-length vinculin tagged to GFP sequence at the 5’ terminus and driven by CAG promoter. **(B)** Cartoon depicting different mutant clones of *vinculin* constructed and tested. Red vertical lines indicate the positions of point mutations.

**Suppl. Figure 3.**
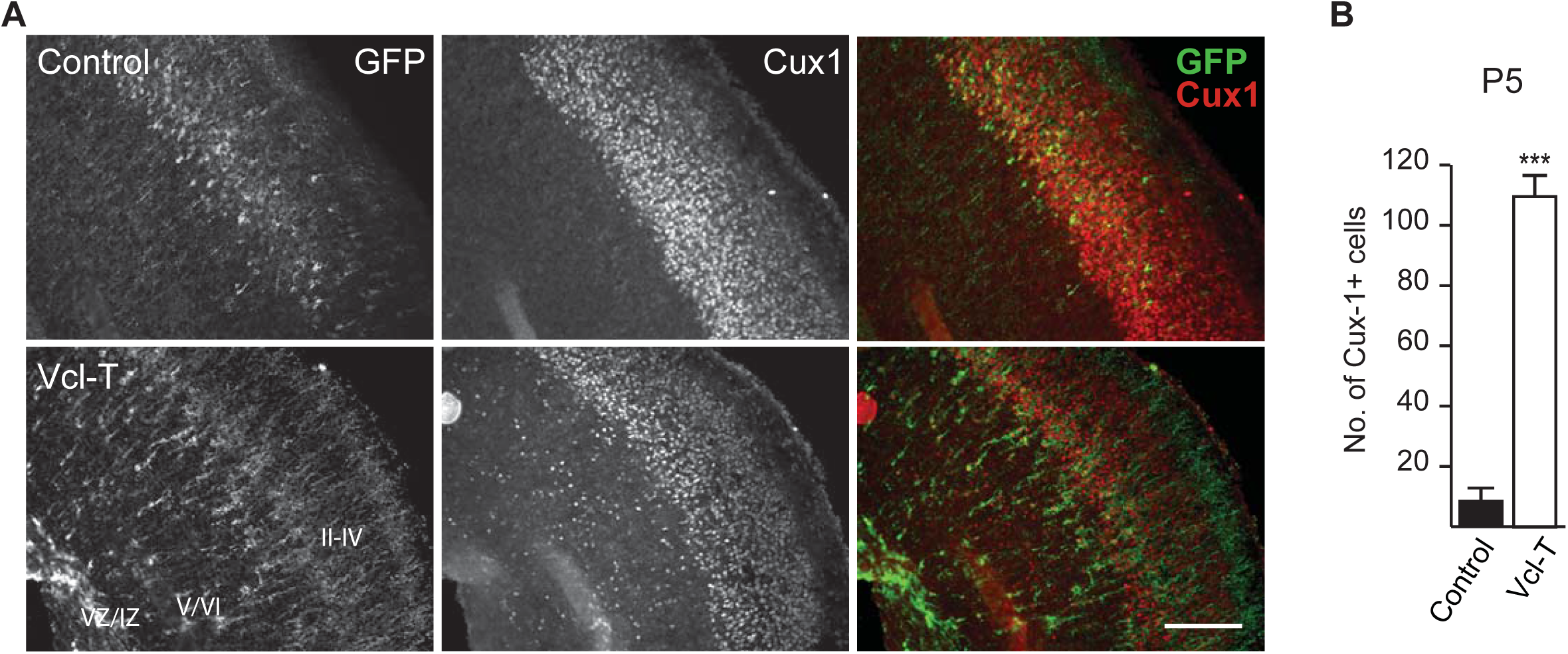
Vcl-T expression affects neocortical lamination. **(A)** Embryos were electroporated at E14.5 with control GFP plasmid or Vcl-T and analysed at P5. 20-µm brains sections were immunostained for GFP and layer 2-4 specific marker, Cux1. In control sections, all Cux1^+^ neurons had reached the upper layers as expected. In contrast, the Vcl-T transfected embryos, many Cux1^+^ neurons were seen occupying the lower layers including around the ventricular zone/intermediate zone (VZ/IZ). Scale, 400 µm. **(B)** Quantification of the number of Cux1^+^ cells in bottom layers of cortex (layers V & VI) and in IZ/VZ from (E) (n=3 animals, \*\*\**p*<0.001. Two-tailed student’s *t* test.

**Suppl. Figure 4.**
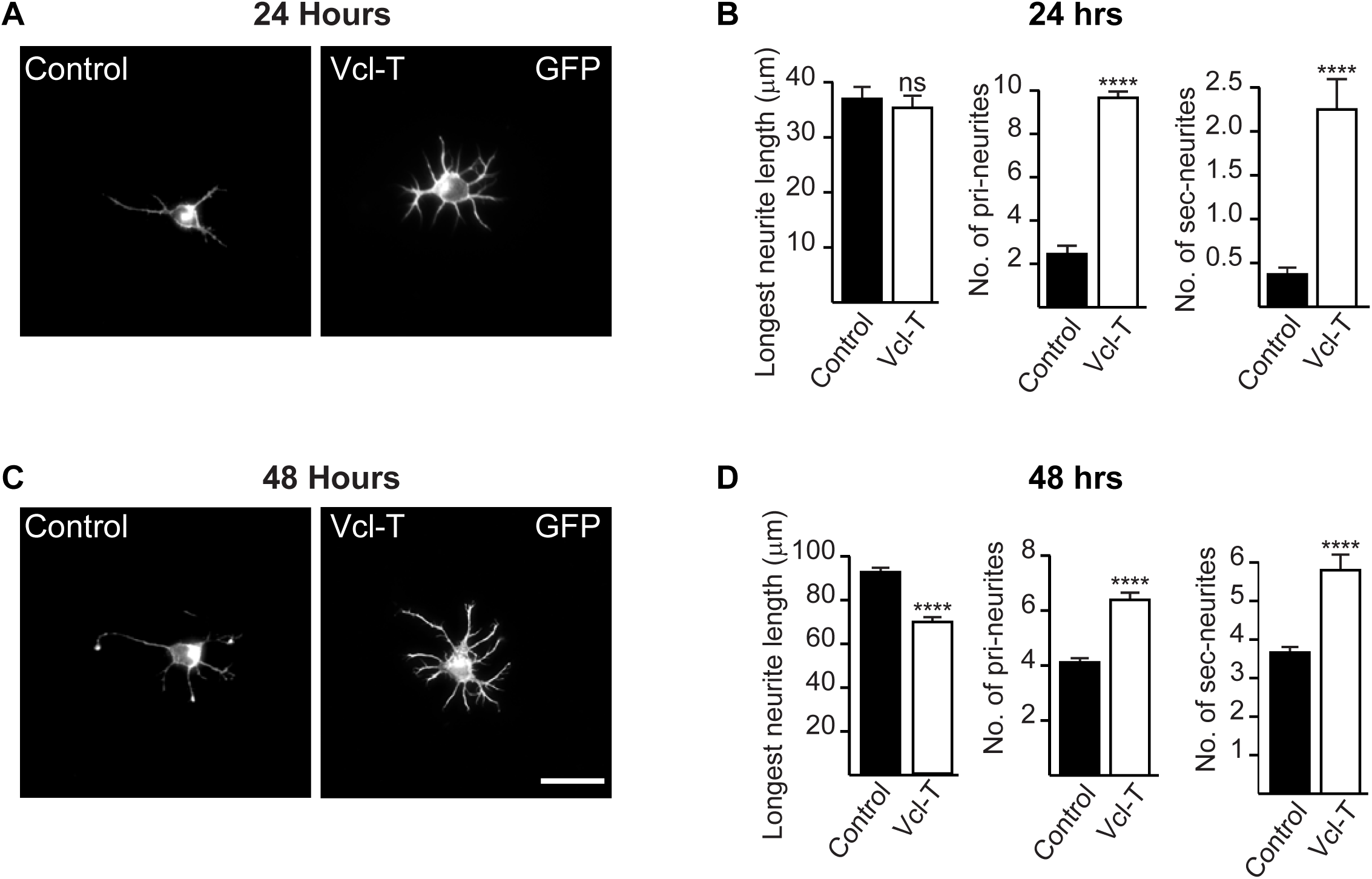
Vcl-T increases branching early in development *in vitro*. Cortical neurons from neonatal mice were transfected with control plasmid and Vcl-T and plated. Cells were fixed at 24 h **(A)** and 48 h **(C)** and visualized for GFP expression. Vcl-T expressing neurons had a greater number of both primary and secondary neurites even at 24 hrs post-plating. Scale, 100µm). (**B, D**) Quantification of the length of longest neurite, number of primary and secondary neurites at 24 hrs **(B)** and 48 hrs **(D)** (n=100 cells, \*\*\*\**p*<0.0001. ns, non-significant). Two-tailed student’s *t* test.

**Suppl. Figure 5.**
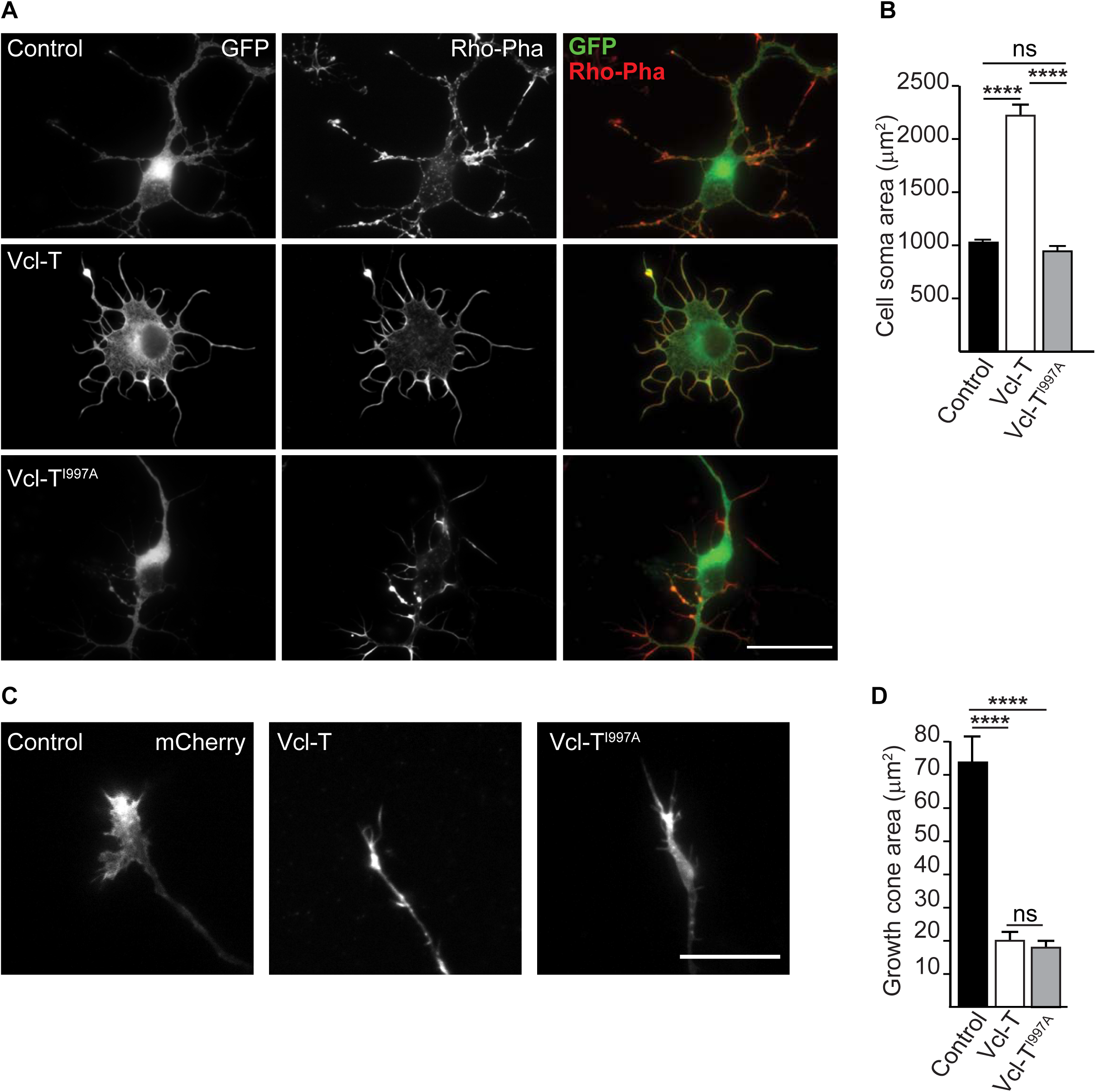
Abolishing actin-Vcl-T interaction reverses cell soma but not growth cone morphology. **(A)** Representative images of neocortical neurons transfected with control vector, Vcl-T and Vcl-T^I997A^ constructs and stained for F-actin using rhodamine-conjugated phalloidin (Rho-Pha). Abolishing interaction between Vcl-T and actin (Vcl-T^I997A^) reduced the cell soma area to that seen in control neurons. **(B)** Quantification of the cell soma area in control, Vcl-T and Vcl-T^I997A^ expressing neurons shown in (A) (n=100 cells). **(C)** Representative images from live imaging of control Vcl-T and Vcl-T^I977A^ expressing neurons co-transfected with LifeAct-mCherry to visualize actin. Abolishing actin-Vcl-T interaction does not rescue growth cone defect exhibited by Vcl-T. **(D)** Quantification of the growth cone area in control, Vcl-T and Vcl-T^I997A^ expressing neurons shown in (C) (n=25 cells). Scale, 50µm (A), 10µm (C). \*\*\*\**p*<0.0001; ns, non-significant. Two-tailed student’s *t* test.

**Suppl. Figure 6.**
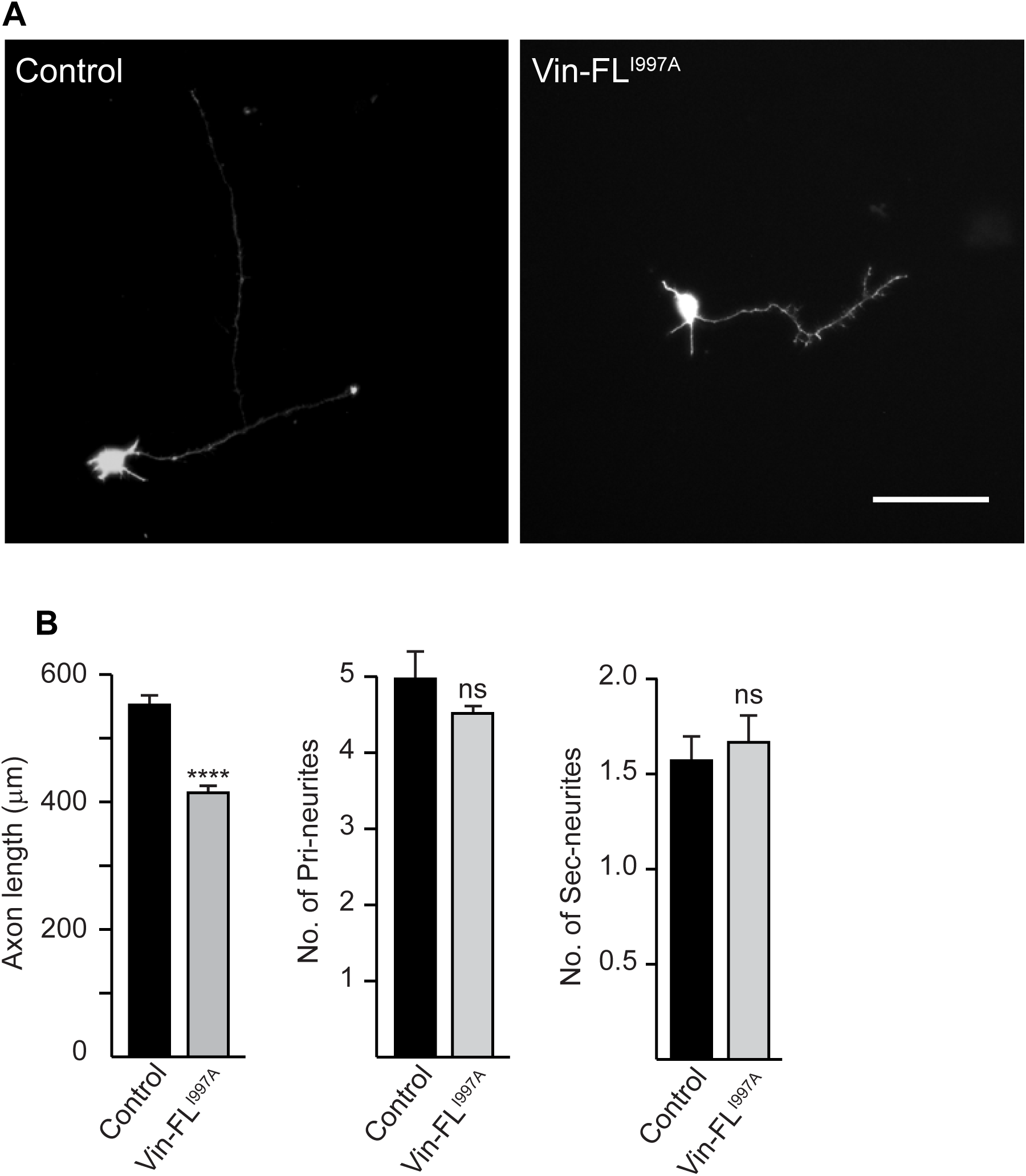
(A) Vinculin full-length defective in actin binding (Vcl-FL^I997A^) does not affect branching. Representative images of neurons electroporated with either empty GFP (control) vector or vinculin full-length defective in actin binding (Vcl-FL^I997A^). Scale, 100µm. **(B)** Quantification of longest neurite length and number of primary and secondary neurites from neurons shown in (A) (n=100 cells). \*\*\*\**p*<0.0001; ns, non-significant. Two-tailed student’s t test.

**Suppl. Figure 7.**
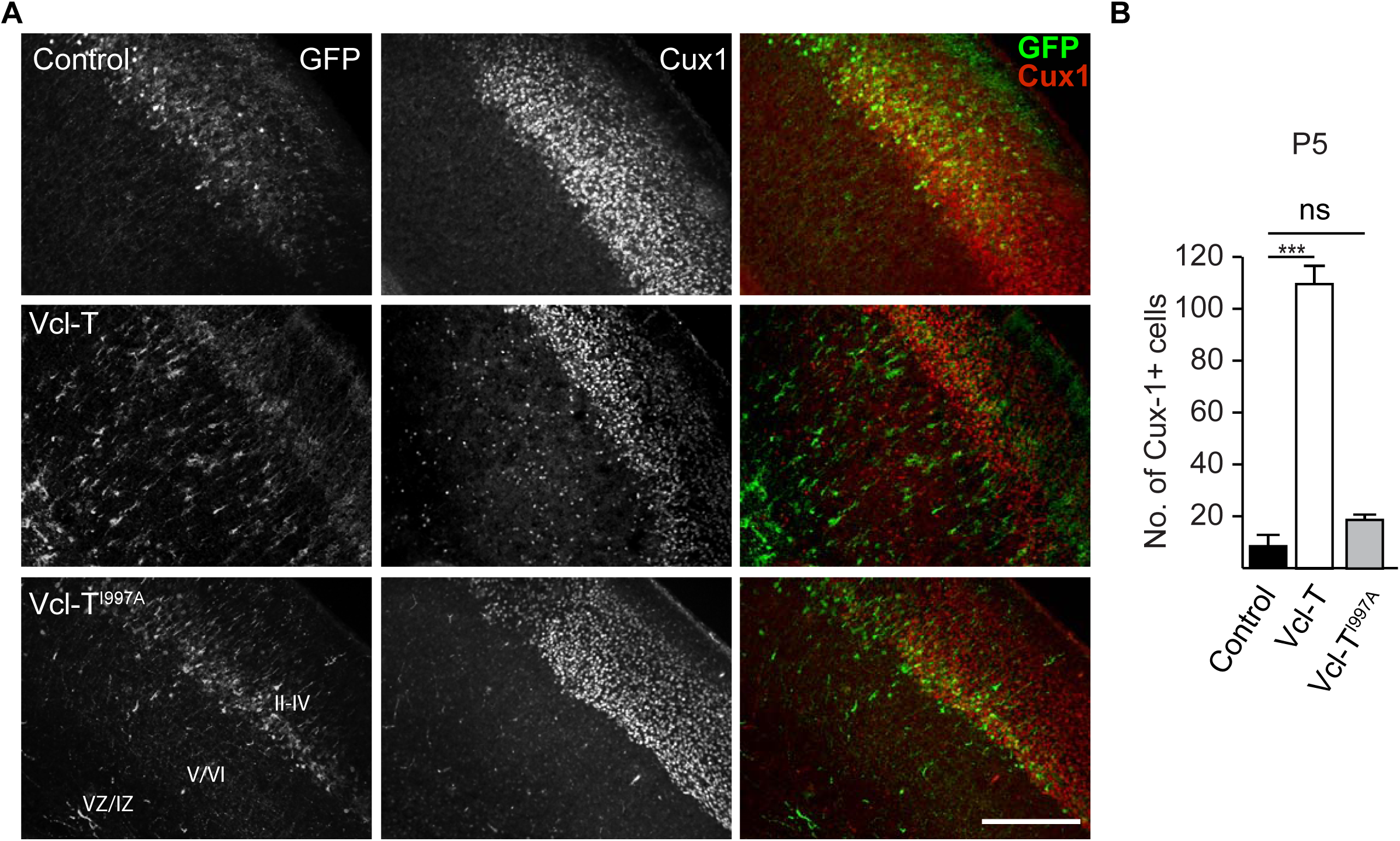
Abolishing actin-Vcl-T interaction rescues cortical lamination defect caused by Vcl-T. **(A)** Analysis of cortical lamination using Cux1 immunostaining shows the rescue of migration of upper layer neurons when actin-Vcl-T interaction is abolished. Scale, 400µm. **(F)** Quantification of the number of Cux1^+^ cells in bottom layers of cortex (layer V & VI) and in IZ/VZ from (A) (n=3 animals) \*\*\**p*<0.001; ns, non-significant. Two-tailed student’s t test.

## Notes

### Competing Interest Statement

The authors have declared no competing interest.

### Summary of Updates

Revised some of the results and the discussion section. Some figure panels in Figures 2 and 5 were replaced.

